# Sinking a giant: quantitative macroevolutionary comparative methods debunk qualitative assumptions

**DOI:** 10.1101/2022.05.05.490811

**Authors:** Matteo Fabbri, Guillermo Navalón, Roger B. J. Benson, Diego Pol, Jingmai O’Connor, Bhart-Anjan S. Bhullar, Gregory M. Erickson, Mark A. Norell, Andrew Orkney, Matthew C. Lamanna, Samir Zouhri, Justine Becker, Cristiano Dal Sasso, Gabriele Bindellini, Simone Maganuco, Marco Auditore, Nizar Ibrahim

**Affiliations:** Negaunee Integrative Research Centre, Field Museum of Natural History, Chicago, IL, USA; Department of Earth Sciences, University of Cambridge, Downing Street, Cambridge, UK; Department of Earth Sciences, University of Oxford, Oxford, UK; Unidad de Paleontología, Departamento de Biología, Universidad Autónoma de Madrid, Madrid, Spain; CONICET, Museo Paleontológico Egidio Feruglio, Trelew, Argentina; Department of Hearth and Planetary Sciences and Peabody Museum of Natural History, Yale University, New Haven, USA; Department of Biological Science, Florida State University, Tallahassee, USA; Division of Vertebrate Paleontology, American Museum of Natural History, New York, NY, USA; Section of Vertebrate Paleontology, Carnegie Museum of Natural History, Pittsburgh, PA, USA; Department of Geology and Health and Environment Laboratory, Hassan II University of Casablanca, Casablanca, Morocco; Department of Biology, University of Detroit Mercy, Detroit, MI, USA; Sezione di Paleontologia dei Vertebrati, Museo di Storia Naturale di Milano, Milan, Italy; Dipartimento di Scienze della Terra ‘A. Desio’, Università degli Studi di Milano, Milan, Italy; Associazione Paleontologica Paleoartistica Italiana, Parma, Italy; School of the Environment, Geography and Geosciences, University of Portsmouth, Portsmouth, UK

## Abstract

Myhrvold et al.^1^ suggest that our inference of subaqueous foraging among spinosaurids^2^ is undermined by selective bone sampling, inadequate statistical procedures, and use of inaccurate ecological categorizations. Myhrvold et al.^1^ ignore major details of our analyses and results, and instead choose to portray our inferences as if they were based on qualitative interpretations of our plots, without providing additional analyses to support their claims. In this manuscript, we thoroughly discuss all the concerns exposed by Myhrvold et al.^1^. Additional analyses based on our original datasets^2^ and novel data presented by Myhrvold et al.^1^ do not change our original interpretations: while the spinosaurid dinosaurs *Spinosaurus* and *Baryonyx* are recovered as subaqueous foragers, *Suchomimus* is inferred as a non-diving animal.

## Main text

Myhrvold et al.^1^ challenge our inference of subaqueous foraging in spinosaurid dinosaurs based on the following three concerns: 1) Accuracy of estimated density of the femoral shafts of *Spinosaurus* and *Suchomimus*, based on CT scan imaging and an additional individual referred to *Spinosaurus*, alongside concerns about air-content in the vertebral column and proportional hindlimb size that could have reduced overall body mass of the neotype of *Spinosaurus*. (2) Concerns about our statistical procedure, including the scope of our comparative dataset and potentially high misclassification rates. (3) That, in their view, we presented a novel redefinition of the term ‘aquatic’.

We closely examine these concerns, supporting the validity of our original bone density measurements^2^ even based on the new scan data presented by Myhrvold et al.^1^ Importantly, we show that the relatively small potential changes to these values do not undermine our hypotheses. *Spinosaurus* (the neotype and the new specimen) and *Baryonyx* have the highest values of femoral compactness of any non-avian dinosaurs yet studied, and also higher than any extant taxa that do not undergo habitual submersion in water, including large-bodied taxa such as rhinos and elephants. Contrary to claims that it is selectively sampled, our comparative dataset^2^ of bone density as an indicator of subaqueous foraging in amniotes is the largest and most phylogenetically comprehensive yet presented. We also demonstrate that our analyses have a very low misclassification rate (which was also reported and discussed in our original paper^2^). Therefore, the ‘analogy’ dataset of human height and sex presented by Myhrvold et al.^1^, which has a very high misclassification rate, is irrelevant to our interpretations. Finally, we make no apology that we defined our ecological variable very clearly prior to analysis, because use of clearly-defined variables is central to valid analysis of ecomorphology and was not intended as a redefinition of the broad concept of what it means to be an aquatic animal.

Before tackling these concerns in detail, we want to specify that the depiction of our methods by Myhrvold et al.^1^ is incorrect and misleading. Our ecological inference^2^ is not based on the visual interpretation of the regression lines and scores in PGLS models or violin plot distribution, as they imply. Instead, our inferences^2^ rest on the use of phylogenetic flexible discriminant analyses (repeated on 100 different trees with variable branch lengths to account for stratigraphic uncertainty) which consider the strong phylogenetic signal in our data (e.g., the average bone density is higher for mammals than for archosaurs). In each iteration, the variables (global bone density, and log-10 transformed midshaft diameter) from the training set of taxa with known ecologies, are used to generate the discriminant functions. Importantly, the discriminant functions are used to re-classify taxa with known ecologies to estimate the accuracy of the method (i.e., misclassification rates). Finally, these validated functions are subsequently used to *a posteriori* predict the ecologies in extinct taxa with unknown ecologies (including spinosaurids), which provides an estimation of species-specific probabilities for each iteration of the analysis. Therefore, our ecological inferences are based on the median probability for each single taxon to be inferred as a subaqueous forager (> 80% cutoff) across 100 iterations, taking into account both the strong phylogenetic structure of bone compactness data and the accuracy of the method as a proxy of the certainty of the ecological inferences. Myhrvold et al.^1^ completely ignore all these details and instead chose to portray our inferences as if they were based on qualitative interpretations of our plots. As such, they mostly based their criticism of our methods on visual interpretations of our plots and perceived inconsistencies in those, without providing additional analyses to support their claims.

### 1) Accuracy of the quantification of skeletal density in spinosaurid dinosaurs, its relationship to mass reduction, and inference of subaqueous foraging in *Spinosaurus* and *Suchomimus* based on novel data

#### Universal definition of density

as every scholar knows, the density of an object is defined as the ratio between its mass and volume. Therefore, mass cannot be assumed to have a linear relationship with density, contrary to the suggestion of Myhrvold et al.^1^. A clear example of this basic physical concept applied to buoyancy is the difference between granite and pumice, which have an average density of 2.7-2.8 g/cm^3^ and 0.5 g/cm^3 3^, respectively. Because the density of water is 1g/ cm^3^, granite sinks in water, contrary to pumice, and regardless of the mass of the object. This also means, that one gram of granite sinks in water, but a 4-ton adult individual of Asian elephant does not (*Elephas maximus*; global bone density = 0.772 (femur) and 0.481 (dorsal rib) in our study^2^). Therefore, Myhrvold et al.^1^’s argument suggesting that body mass reduction in *Spinosaurus* due to relative shortening of the hindlimbs has implications for its buoyancy control is ill-founded.

#### Skeletal distribution of osteosclerosis and pneumatization

Myhrvold et al.^1^ state that axial pneumatization is present along the cervical and caudal axial skeleton of *Spinosaurus* with consequences for their buoyancy control. However, no pneumatization is observed in the caudal vertebrae of *Spinosaurus* (Figure 1), only fragments of a proximal caudal vertebra are known in *Bayronyx*^4^, and proximal caudals are not known in *Suchomimus* (Figure 1 in our original study^2^). The lack of a volumetric quantification of density reduction caused by vertebral pneumatization in spinosaurids makes the argument of Myhrvold et al.^1^ purely qualitative, and untestable based on the currently available data. We note that spinosaurids have a reduced amount of vertebral pneumatization in the trunk and sacrum, compared to many other multi-tonne theropods^5^. Furthermore, high bone density is present across the entire postcranial skeleton in *Spinosaurus* (global bone density > 0.9 for: dorsal ribs, dorsal and caudal neural spines, femur, tibia, and fibula) and *Baryonyx* (global bone compactness > 0.87 for: dorsal ribs, scapula, pubis, ischium, femur and fibula), as stated and quantified in our original study^2^ (Figure 1 and Extended Data Figure 10), and also including the manual phalanx of the neotype specimen of *Spinosaurus* (Figure 1) which completely lacks a medullary cavity.

**Figure 1.**
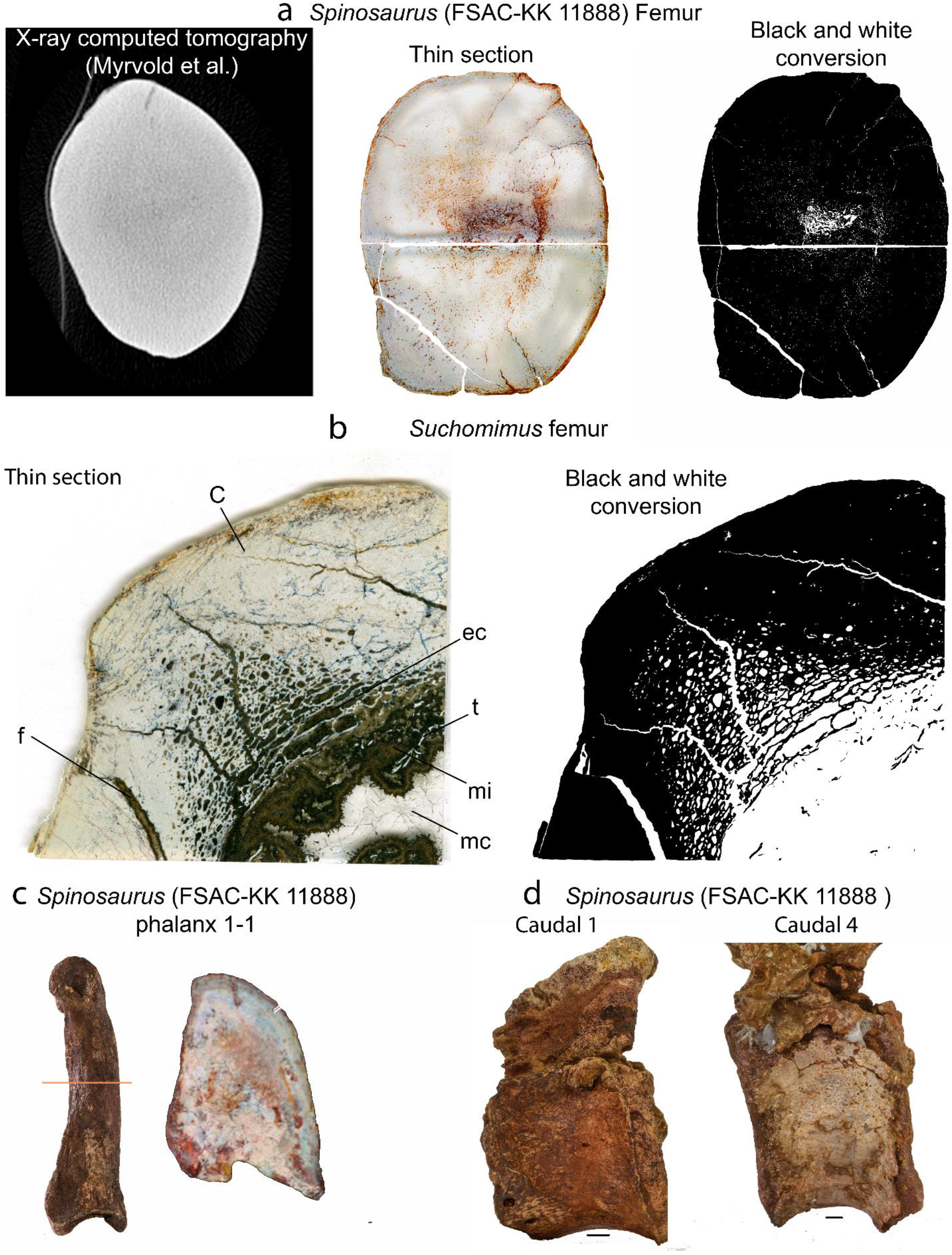
a) Comparison of data sources for the femur of the neotypic specimen of *Spinosaurus*: contrary to our thin section of this element, the CT scan data reported by Myhrvold et al.^1^ lack any kind of contrast, obscuring any internal structure and bone tissue distribution; b) close up of the femoral thin section of *Suchomimus*: Myhrvold et al.^1^ suggest that the bone density referred to this specimen was underestimated in our study, due to their erroneous interpretation of sediment and mineral infilling of the medullary cavity as bone tissue; c) based on CT scan data, Myhrvold et al.^1^ state that a single phalanx of the neotype of *Spinosaurus* possess a medullary cavity, invalidating our inference of widespread osteosclerosis across the postcranium of this animal; we show here that a cross section of the phalanx lacks any medullary cavity, as previously described in Ibrahim et al.^13–14^; d) Caudal vertebrae 1 and 4 of the neotype of *Spinosaurus:* contrary to what suggested by Myhrvold et al.^1^, no pneumatization is present in the caudal region of this taxon. Abbreviations: c=cortex; ec=erosional cavity; f=fracture; mc=medullary cavity; mi=mineral infilling; t=trabeculae. Scale bar in (d) is 10 mm.

Ironically, Myhrvold et al.^1^ suggest that *Spinosaurus* could be more likely interpreted as a terrestrial animal rather than a subaqueous forager based on a single qualitative trait (tibia/femur length ratio=> 1), while emphasizing the impossibility of ecological inference based on quantitative methods when categories overlap, without regard to the fact that misclassification rates are low for our analyses. The problems with their reasoning are clear given that all modern birds, including subaqueous foragers, are characterized by a tibia/femur length ratio of more than 1, including penguins^e.g.6^.

Myhrvold et al.^1^ suggest that the sections used for the three spinosaurid taxa analyzed in our study^2^ come from distal portions of the femur, implying that we are underestimating the bone density in these species (assuming that bone density is highest in the midshaft). Assuming that they are correct, this does not explain variation in bone density among spinosaurids — *Baryonyx* and *Spinosaurus* show some of the highest bone density values observed among amniotes, while *Suchomimus* is more comparable to other small and large predatory dinosaurs.

Myhrvold et al.^1^ suggest that we underestimated bone density in *Suchomimus* during the conversion of the femoral thin section into a black & white figure (the curating step prior to estimation of bone compactness), causing us to mis-identify bone as rock matrix. However, we did not apply our techniques blindly, but instead used careful observation to quantify bone compactness. As shown in Figure 1, the bone tissue in this specimen has a distinct white hue: Myhrvold et al.^1^ conflate the mineral infilling surrounding the trabecular bone and bone tissue. Additionally, based on CT scan imaging, Myhrvold et al.^1^ accuse us of ignoring a medullary cavity in the femur of the neotypic specimen of *Spinosaurus* and that we are incorrectly oversampling bone tissue based on a thin section of the femur. As shown in Figure 1, cross sections obtained from the CT scan presented by Myhrvold et al.^1^ lack adequate contrast and resolution, obscuring any details of its internal structure, contrary to the thin section used in our study^2^ (Figure 1). Surprisingly, Myhrvold et al.^1^ do not comment on our results^2^ based on the dorsal ribs of *Spinosaurus* and *Baryonyx*, which still recover these two taxa as subaqueous foragers with high probabilities.

Myhrvold et al.^1^ describe a different individual of *Spinosaurus* (partial, isolated femur) characterized by an open medullary cavity, potentially invalidating our inference of subaqueous foraging. We consider important to stress that many modern diving species exhibit an open medullary cavity, contrary to what Myhrvold et al.^1^ assume (e.g., *Caiman* in Figure 1 in our original study^2^, penguins, sauropterygians, the hippopotamus, and others in Extended data figures 1-7 in our original study^2^). In order to see if this isolated femur of *Spinosaurus* challenges our inference of its ecology, we translated the cross section of this specimen into black & white and quantified its bone density in the software BoneProfiler^7^, which is recovered as 0.941: this is slightly lower than that of the neotype (bd=0.968), but is still remarkably high among amniotes. This bone density value was substituted to the one of the neotype in our original dataset to infer ecological behaviour in this new individual, following the methods of our study. Similarly, we also substituted our previous value of bone density for the femur of *Suchomimus* with a novel estimate obtained from a cross section extrapolated from the low-quality CT scan data presented by Myhrvold et al.^1^: the section we used come from an area closer to the fourth trochanter (sections 4-3 in Supplementary Figure 1 in Myhrvold et al.^1^). The resulting bone density is only slightly higher (bd=0.756) than our original value for *Suchomimus* and might be related with the low resolution of the scan data. Notwithstanding the inadequate quality of the imaging of the new specimens, our ecological inference based on phylogenetic discriminant analyses for these taxa does not change, giving similar probabilities for *Spinosaurus* and *Suchomimus* as subaqueous forager (now based on two separate individuals) and non-diving, respectively (see Table 1), and demonstrates how our ecological inference is independent from the sampling region of the femora.

**Table 1.**
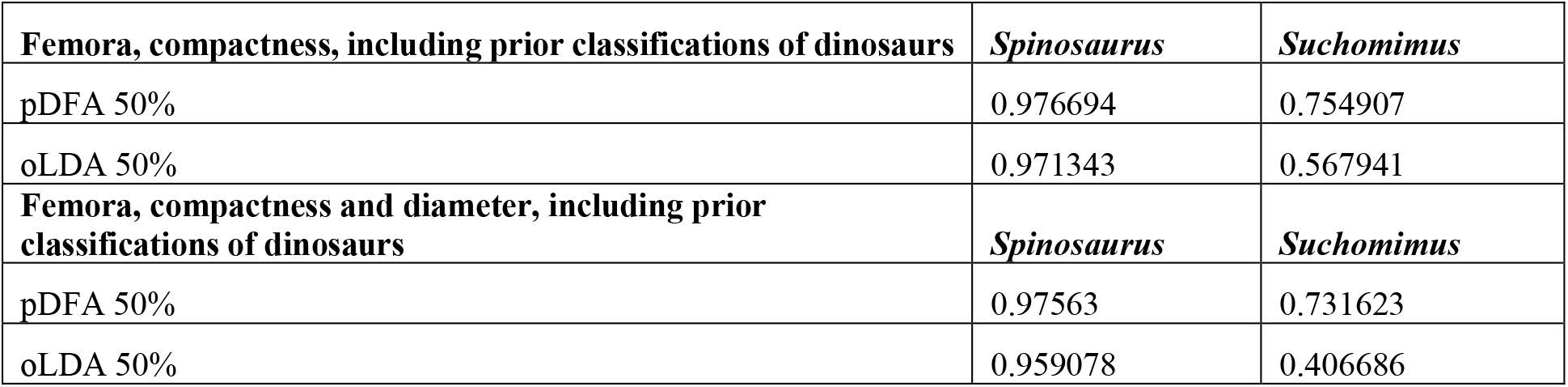
Posterior probability of subaqueous foraging for the novel individual of *Spinosaurus* and additional section of *Suchomimus* by Myhrvold et al.^1^. *Spinosaurus* is still recovered as a subaqueous forager, contrary to *Suchomimus* which is inferred as a non-diver.

| **Femora, compactness, including prior classifications of dinosaurs** | *Spinosaurus* | *Suchomimus* |
| --- | --- | --- |
| pDFA 50% | 0.976694 | 0.754907 |
| oLDA 50% | 0.971343 | 0.567941 |
| **Femora, compactness and diameter, including prior classifications of dinosaurs** | *Spinosaurus* | *Suchomimus* |
| pDFA 50% | 0.97563 | 0.731623 |
| oLDA 50% | 0.959078 | 0.406686 |

### 2) Statistical procedure: comparative dataset, phylogenetic correction, and classification rates

#### The impact of higher bone density among extinct subaqueous taxa

Myhrvold et al.^1^ state that the highest bone density values in our dataset are represented by extinct taxa, and that this undermines the predictive power of ecological behavior in extinct taxa because it is not based on neontological observations. Although the 15 highest bone density values in the femur dataset belong to extinct stem cetaceans, sauropterygians, and early seals, the difference in bone density between *Serpianosaurus* (an early sauropterygian and the densest amniote in our dataset^2^) and *Caiman* (the densest extant taxon in our study^2^) is only 0.06. As shown in Figure 2, there is a clear overlap between the range of bone density values (both femur and dorsal ribs) characterizing extant and extinct diving animals. A possible explanation for extant species exhibiting a broader range of bone compactness values than their extinct counterparts is that we only scored the latter as ‘subaqueous foragers’ when the adaptations for this behaviour are uncontested. In contrast, extant taxa that are anatomically less specialized for subaqueous foraging (potentially with lower bone densities) could be classified as such based on observation of their in-life behaviour. Therefore, unless Myhrvold et al.^1^ is suggesting that stem whales, such as remingtonocetids, basilosaurids, and protocetids, and sauropterygians, including plesiosaurs, were not subaqueous foragers, we don’t see how this challenges our results. Furthermore, some of the extinct taxa that we include are crown members of recent clades of aquatic animals like phocids or otariids or closely related stem members of recent groups with similar ecologies (e.g. penguins). The values between these fossils and their extant counterparts are very similar suggesting there is no taphonomic bias in our quantification of bone density in extinct taxa, contrary to the suggestions by Myhrvold et al.^1^.

**Figure 2.**
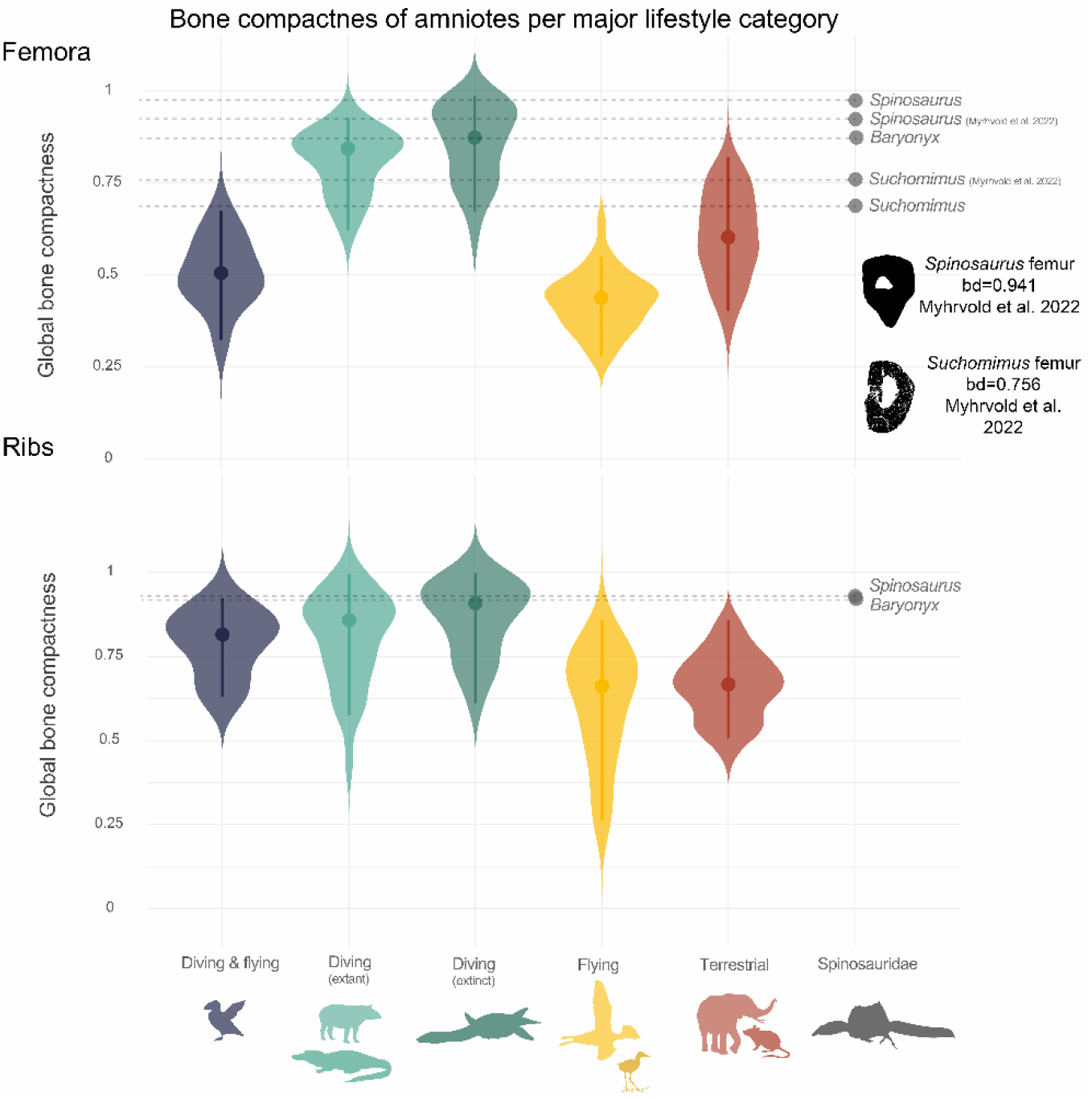
Violin plots separating diving extant and extinct taxa included in our femoral and dorsal rib datasets: the values between extinct and extant divers broadly overlap between each other.

#### Lack of large modern divers

according to Myhrvold et al.^1^, the absence of large, extant diving animals comparable to *Spinosaurus* in body size does not allow us to properly discern between the influence of allometry and ecological adaptations. Although the femur dataset does not contain modern, large cetaceans because of extreme reduction/loss of hindlimbs in this clade, unequivocal subaqueous extinct taxa of large size, such as basilosaurids (early whales), mosasaurs, and plesiosaurs, do have comparable or even larger body size than *Spinosaurus*. Our rib dataset^2^ indeed contains many large size extant taxa including large living cetaceans (e.g. *Balaenoptera* and *Orcinus*) or living sirenians with very wide ribs (e.g., *Trichechus*) along with large and unequivocally aquatic large sized extinct reptiles such as mosasaurs (e.g. *Tylosaurus*), taxa that exceed the body size of *Spinosaurus*. The bone density of dorsal ribs and femora of *Spinosaurus* and *Baryonyx* are comparable or exceed those seen in all these clades, consistent with the inference of subaqueous foraging in *Spinosaurus* and *Baryonyx*.

#### The role of allometry and graviportality

Myhrvold et al.^1^ state that high bone density in *Spinosaurus* and *Baryonyx* is the result of allometry, rather than ecology, based on a qualitative assessment of the vicinity of data points in our PGLS and violin plots (“*Loxodonta* falls close to *Baryonyx*”), their perceived deficit of terrestrial large animals (“the sauropod *Alamosaurus*, the ornithischian *Stegosaurus*, and the African elephant *Loxodonta*” in the femur dataset), and the correlation between density and cross-sectional diameter in a subsample of six nothosaurs.

We already quantified the role of allometry as an explanatory variable for variation of bone density in our dataset^2^, comparing this to other multiple regression models using the corrected Akaike information criterion (AICc). Femoral midshaft diameter is strongly correlated to body mass in amniotes^e.g.8^ and we used this as a proxy for overall size. Adding this proxy to a regression model “bone density ~ subaqueous foraging” (AICc weight = 0.673 [femur], 0.638 [ribs]) results in a model “bone density ~ subaqueous foraging + body mass” results in a substantially worse explanation of variation in bone density (AICc weight = 0.014 [femur], 0.019 [ribs]), and the coefficient of our body mass proxy is widely non-significant (Supplementary Tables 3-4 in our study^2^). This rejects the hypothesis that bone density shows a strong allometric influence. Instead, the model “bone density-subaqueous foraging” is found to be the best explanatory model on both the datasets^2^ (femur and ribs).

Variation in body size is therefore, at best, a minor explanatory factor for variation in bone density in amniotes, as we discussed in our original paper^2^. Nevertheless, our original phylogenetic discriminant analyses^2^ included both global compactness and log-10 transformed midshaft diameter to establish predictions of individual taxa. Therefore, our original analyses^2^ fully evaluated the hypothesis that allometry is an important factor in explaining variation in bone density and our discriminant analyses take variation in size into account in order to establish predictions. The example of six different species of *Nothosaurus* in support of the argument of allometry as a driving factor of bone density by Myhrvold et al.^1^ is “cherry-picking”, and may be misleading given that our analyses are based on comparative analysis of hundreds of taxa across the amniote phylogeny.

Myhrvold et al.^1^ also ignore the presence of two sauropod dinosaurs in our dataset, namely *Alamosaurus* and *Antetonitrus*, as well as large predatory dinosaurs of larger size than the neotype of *Spinosaurus* (cross-sectional diameter=81.52 mm), namely *Tyrannosaurus* (197 mm), *Baryonyx* (154 mm), *Suchomimus* (120.6 mm), *Torvosaurus* (132.57), *Allosaurus* (89.49), and *Asfaltovenator* (103.03 mm), in addition to *Loxodonta* and other large modern mammals in the femur dataset. Therefore, a comparative framework of large terrestrial animals is present; yet, *Spinosaurus* (bd=0.968) and *Baryonyx* (bd=0.876) show remarkably higher bone density for the femur than any other large terrestrial amniote (*Alamosaurus* (bd=0.777), *Loxodonta* (bd=0.855), *Stegosaurus* (bd=0.81), and the other large theropods (highest and lowest values ranging from bd=0.47-0.728 in *Asfaltovenator* and *Torvosaurus*, respectively). Interestingly, Myhrvold et al.^1^ decided not to comment on our rib dataset, which includes five large sauropods (*Alamosaurus*, *Brachiosaurus*, *Apatosaurus*, *Diplodocus*, and *Spinophorosaurus*), *Mammuthus* and modern elephants, a large carcharodontosaurid, and modern cetaceans (e.g., *Balaenoptera* or *Orcinus*):

#### The influence of allometry and mixed ecologies for inferring ecology in extinct taxa

according to the qualitative concerns proposed by Myhrvold et al.^1^, our inference of subaqueous foraging in extinct taxa is faulted by the inclusion of modern taxa capable of both flight and diving, the influence of allometry in our predictive model, and the lack of categorization of some extinct taxa with unquestioned ecologies such as large terrestrial non avian dinosaurs (which we scored as unknown in our original study). In order to test the influence of these factors on our probabilistic inference of subaqueous foraging among non-avian dinosaurs, we ran a new batch of pDFA analyses using both the femur and the dorsal rib datasets including (as in the original paper) and excluding midshaft diameter. To facilitate interpretation of misclassification rates, we ran these analyses over 1000 iterations, each time re-sampling our dataset to equal counts of taxa in each class (i.e. equal counts of subaqueous foragers and non-subaqueous foragers), also selecting a different tree from our phylogenetic distribution each time to reflect stratigraphic uncertainty. We varied two aspects of the inputs to our analysis: (i) We included and excluded prior classifications of some non-avian dinosaurs in the training set; (ii) We include and excluded information on body size, based on our body size proxy, femoral and dorsal rib midshaft diameter. Graviportal and pelagic taxa were included in all analyses. These analyses return a correct classification rate of 82-85% (femora) or 74-79% (ribs) for taxa in the training set using phylogenetic flexible discriminant function analysis, or 84–85% (femora) and 75–82% (ribs) for ordinary (non-phylogenetic) linear discriminant analysis. These results also demonstrate that our analyses in general have low misclassification rates, and that allometry and prior categorizations of non-avian dinosaurs have only a minimal impact on performance of our discriminant analyses. The posterior probabilities from these analyses recover *Spinosaurus* and *Baryonyx* as subaqueous foragers, while *Suchomimus* is recovered as a non-diving species (Table 3).

**Table 2.** Correct classification rates (i) including and excluding prior classifications of some non-avian dinosaurs in the training set, and (ii) including and excluding information on body size using both the femur and the dorsal rib datasets. Correct classification rate remains high and in the same ranges of our original study^2^.

| **Model** | **pDFA:** | **oLDA:** |
| --- | --- | --- |
| **femur_compactness excluding prior classification of other non-avian dinosaurs** | 0.84840678 | 0.844610169 |
| **femur_compactness including prior classifications of dinosaurs** | 0.820127119 | 0.850864407 |
| **femur_compactness and diameter excluding prior classification of other non-avian dinosaurs** | 0.85590678 | 0.849186441 |
| **femur_compactness and diameter including prior classifications of dinosaurs** | 0.820508475 | 0.854771186 |
| **rib_compactness excluding prior classification of other non-avian dinosaurs** | 0.748061224 | 0.773704082 |
| **rib_compactness including prior classifications of dinosaurs** | 0.741581633 | 0.756173469 |
| **rib_compactness excluding prior classification of other non-avian dinosaurs** | 0.792408163 | 0.822163265 |
| **rib_compactness and diameter including prior classifications of dinosaurs** | 0.775438776 | 0.800020408 |

**Table 3.**
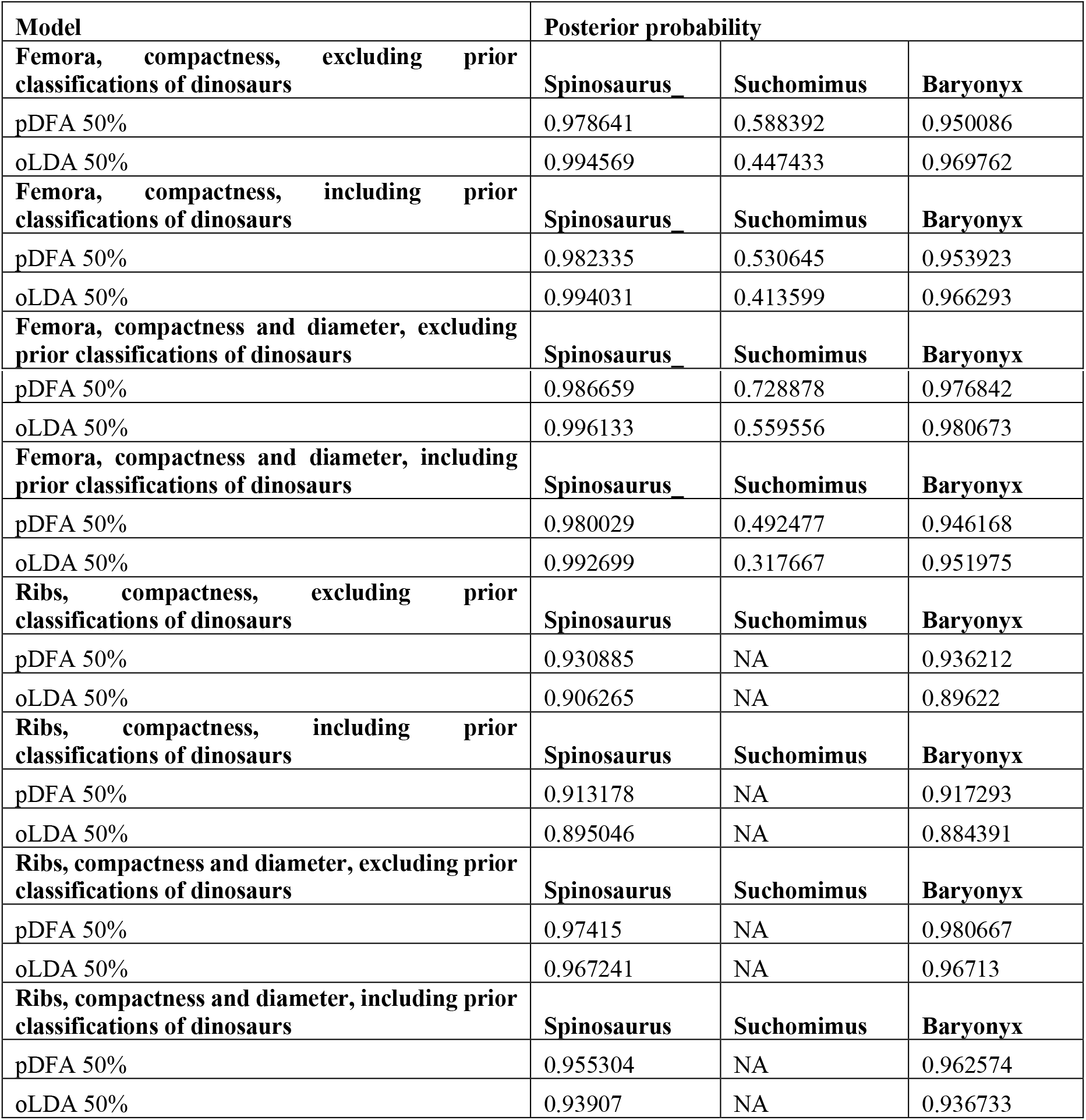
Posterior probability of subaqueous foraging in *Spinosaurus*, *Suchomimus* and *Baryonyx* based on the novel analysis presented in this study (i) including and excluding prior classifications of some non-avian dinosaurs in the training set, and (ii) including and excluding information on body size using both the femur and the dorsal rib datasets. *Spinosaurus* and *Baryonyx* are recovered as subaqueous foragers, while *Suchomimus* is inferred as anon-diver.

| **Model** | **Posterior probability** |  |  |
| --- | --- | --- | --- |
| **Femora, compactness, excluding prior classifications of dinosaurs** | **Spinosaurus_** | **Suchomimus** | **Baryonyx** |
| pDFA 50% | 0.978641 | 0.588392 | 0.950086 |
| oLDA 50% | 0.994569 | 0.447433 | 0.969762 |
| **Femora, compactness, including prior classifications of dinosaurs** | **Spinosaurus_** | **Suchomimus** | **Baryonyx** |
| pDFA 50% | 0.982335 | 0.530645 | 0.953923 |
| oLDA 50% | 0.994031 | 0.413599 | 0.966293 |
| **Femora, compactness and diameter, excluding prior classifications of dinosaurs** | **Spinosaurus_** | **Suchomimus** | **Baryonyx** |
| pDFA 50% | 0.986659 | 0.728878 | 0.976842 |
| oLDA 50% | 0.996133 | 0.559556 | 0.980673 |
| **Femora, compactness and diameter, including prior classifications of dinosaurs** | **Spinosaurus** | **Suchomimus** | **Baryonyx** |
| pDFA 50% | 0.980029 | 0.492477 | 0.946168 |
| oLDA 50% | 0.992699 | 0.317667 | 0.951975 |
| **Ribs, compactness, excluding prior classifications of dinosaurs** | **Spinosaurus** | **Suchomimus** | **Baryonyx** |
| pDFA 50% | 0.930885 | NA | 0.936212 |
| oLDA 50% | 0.906265 | NA | 0.89622 |
| **Ribs, compactness, including prior classifications of dinosaurs** | **Spinosaurus** | **Suchomimus** | **Baryonyx** |
| pDFA 50% | 0.913178 | NA | 0.917293 |
| oLDA 50% | 0.895046 | NA | 0.884391 |
| **Ribs, compactness and diameter, excluding prior classifications of dinosaurs** | **Spinosaurus** | **Suchomimus** | **Baryonyx** |
| pDFA 50% | 0.97415 | NA | 0.980667 |
| oLDA 50% | 0.967241 | NA | 0.96713 |
| **Ribs, compactness and diameter, including prior classifications of dinosaurs** | **Spinosaurus** | **Suchomimus** | **Baryonyx** |
| pDFA 50% | 0.955304 | NA | 0.962574 |
| oLDA 50% | 0.93907 | NA | 0.936733 |

#### The “ecological fallacy” and the impossibility to reconstruct ecological adaptations in extinct taxa

the ecological fallacy^9^ formalizes how the behaviour of groups/populations might not be representative of single individuals. Myhrvold et al.^1^ invoke the “the ecological fallacy” to suggest that individual predictions of subaqueous ecology cannot be derived from bone density group distributions and present an example of height distribution per sex in humans to prove their point. This argument boldly negates the possibility of ecomorphological studies but ignores that the fact that sex cannot be confidently inferred from height data using that dataset of human heights is due to a very strong overlapping in the sex-specific distributions resulting in a high misclassification rate as stated before, bone density in our data exhibits a low misclassification rate of species that engage in habitual submersion. As a result, the analogy to a human height/sex dataset with high misclassification rate presented by Myhrvold et al.^1^ is not appropriate. Our analyses^2^ take into account phylogenetic relationships and varying branches, to draw posteriori probabilistic inferences of ecological adaptations in extinct taxa. We consider this approach superior to conclusions solely based on group distributions and/or proximity to the regression lines of any ecological group. Myhrvold et al.^1^ seem to claim that this is our primary approach for the inference of ecological behavior in these two taxa, which is completely incorrect. Ironically, the suggestion in Myhrvold et al.^1^ that allometry influences bone density such that *Baryonyx* and *Spinosaurus* might be rather regarded as graviportal animals, is driven by their assumption that the distribution of taxa in our PGLS (group distribution and/or proximity to the regression line) is directly informative for inferring ecological behavior in extinct taxa. Qualitative assessments presented by Myhrvold et al.^1^ challenging our results (e.g. “*Loxodonta* falls close to *Baryonyx*” and “*Phoca* is closer to the “flying” regression line”) assume that a couple of data points can be used to generalized certain trends within groups and vice versa, exactly falling for what the “ecological fallacy” describes^9^.

### 3) Definition of subaqueous foraging and its application to previous autoecological categorizations

As we explained in our manuscript^2^, our ecomorphological attribution is focused on statistical tests of a specific behaviour linked to an ecology and to bone density, rather than representing an attempt to describe semi-aquatic or aquatic ecologies in their entirety. We find our categorization to be more precise: for example, previous studies^e.g.10^ coded penguins and cetaceans as ‘aquatic’, while crocodilians were stated as ‘semiaquatic’. Whereas penguins and crocodilians are still ecologically dependent on terrestrial environments (for example, for laying eggs) and cetaceans are not, all these animals engage in habitual subaquatic submersion and frequent subaqueous foraging. Therefore, our ecological attribution is in agreement with previously applied ecological categories ^e.g.10^, but do not exclude dependency to terrestrial environments to satisfy autecological requirements, such as reproductive behaviour. Although habitual submersion, as epitomized by the frequent use of subaqueous foraging, is only one functionally important aspect of aquatic behaviour, it is the key aspect that we hypothesized as having a functional relationship to bone density. That hypothesis is supported by our analyses^2^, whereas other aspects of aquatic ecologies are not. Although we believe that spinosaurids were not independent from terrestrial environments (e.g. for laying eggs^11^), our data and results based on comparative methods give a clear answer regarding their capabilities for subaqueous foraging.

Furthermore, although hippopotamuses, beavers or tapirs do not often forage underwater, as Myhrvold et al.^1^ correctly pointed out, these animals engage in habitual submersion for other purposes such as concealment or refuge, as discussed in our original paper^2^. These behaviours still represent the same biomechanical challenges and the discussion here is terminological. We acknowledge that we could have used another term such as ‘subaqueous submersion’ to describe, what in essence, is the same behaviour. Also, it is important to point out that these exceptions are strictly related to a specific diet: herbivory. Since Stromer 1915^12^, fishing habits have been suggested among spinosaurids. Therefore, unless Myhrvold et al.^1^ is suggesting that spinosaurids might have been herbivores (or would find fish in land), our inference of subaqueous foraging (or habitual submersion) in spinosaurs still stand.

Finally, as we extensively commented on our published manuscript^2^, our results never excluded wading behavior in extinct taxa: our ecological inference based on bone density only allowed us to discern between subaqueous foraging or not. Contrary to what Myhrvold et al.^1^ report, we never stated in our manuscript^2^ that *Suchomimus* was “fully terrestrial”, but we described this taxon as a non-diver.

## Conclusions

In conclusion, we proposed inference of ecological behavior among non-avian dinosaurs based on the largest and most phylogenetically inclusive dataset of bone density ever assembled. We carefully examined all the concerns exposed in Myhrvold et al.^1^. Our results are still confirmed. The reason for this is because poor quality data and opinionated arguments will never be more accurate than appropriate quantitative macroevolutionary comparative studies.

## Notes

### Competing Interest Statement

The authors have declared no competing interest.

